# Inferring polygenic negative selection underlying an individual trait as a distribution of fitness effects (DFEs) from GWAS summary statistics

**DOI:** 10.1101/2024.07.29.601707

**Authors:** Alexander T. Xue, Yi-Fei Huang, Adam Siepel

## Abstract

There has been rising interest in exploiting data from genome-wide association studies (GWAS) to detect a genetic signature of natural selection acting on a given phenotype. However, current approaches are unable to directly estimate the distribution of fitness effects (DFE), an established property in population genetics that can elucidate genomic architecture pertaining to a particular focal trait. To this end, we introduce ASSESS, an inferential method that exploits the Poisson Random Field (PRF) to model selection coefficients from genome-wide allele count data, while jointly conditioning GWAS summary statistics on a latent distribution of phenotypic effect sizes. This probabilistic model is unified under the assumption of an explicit relationship between fitness and trait effect to yield a DFE. To gauge the performance of ASSESS, we enlisted various simulation experiments that covered a range of usage cases and model misspecifications, which revealed accurate recovery of the underlying selection signal. As a further proof-of-concept, ASSESS was applied to an array of publicly available human trait data, whereby we replicated previously published empirical findings from an alternative methodology. These demonstrations illustrate the potential of ASSESS to satisfy an increasing need for powerful yet convenient population genomic inference from GWAS summary statistics.

**Author Summary:** The growth of genome-wide association studies (GWAS) over the past decade has provided a wealth of resources for uncovering the genomic architecture underlying complex traits, including the footprint of selection. Currently, there are computational tools for inferring natural selection whereby GWAS results are leveraged to conduct a binary test for overall presence, estimate a correlated property, or summarize polygenic selection strength with a single statistic. However, a methodology that exploits GWAS data to estimate the distribution of fitness effects (DFE), which is the most direct measurement for the genetic impact of natural selection acting on a complex trait, does not currently exist. To this end, we constructed an approach to directly infer the DFE, wherein per-site selection coefficients specifically associated with a focal trait are aggregated across the genome. This implementation is designed to explicitly model an entire genome-wide set of summary statistics output from a GWAS rather than the individual-level input data, which offers computational efficiency and convenience as well as alleviates privacy concerns. We expect this to be a promising development given the further accumulation of GWAS results and investigators seeking more sophisticated analyses into the relationship between genetics and traits.

## Introduction

A central focus of human genetics is to elucidate the genomic foundation of complex traits. Genome-wide association studies (GWAS), which deploy a regression analysis that maps phenotypes against genotypes, have been a long-standing approach to accomplish this task. Conducting a GWAS typically produces summary statistics for each genetic site, such as an estimated effect size of the genetic variant on the trait of interest as well as an associated standard error in this estimated value [1]. A GWAS typically aims to reveal key genetic contributors by isolating loci with large estimated effect sizes and relatively low standard error, yet most traits are found to be highly polygenic with predominantly small effect sizes. While such results do not quite fulfill the aspirations initially intended when GWAS were first performed over a decade ago, there is still much information contained within the data that can be exploited to gain broader knowledge about genomic processes [2,3]. Therefore, GWAS research has shifted towards developing alternative and downstream methods that consider the full set of variant associations with a focal trait to address questions of genomic architecture, population genetics, and evolutionary ecology [4,5].

One application that is of widespread interest is to utilize allele sample frequencies of single nucleotide polymorphisms (SNPs) to unveil a signature of selection underpinning a polygenic trait [6–16]. Currently, available tools are designed for binary classification of overall presence versus absence, or indirect quantification of genome-wide fitness by parameterizing a proxy property such as the correlation between allele frequency and true effect size. However, a desirable alternative would be to instead directly estimate the distribution of fitness effects (DFE), which is a frequency histogram that consolidates locus-specific selection coefficients throughout the genome [5,17]. As a fundamental concept in evolutionary genetics, the DFE borrows from a long-standing theoretical basis to allow a clearer understanding of population dynamics. Specifically, it is a composite that reflects magnitude and direction of selection, genomic architecture, mutation rates and patterns, demographic history, and other molecular ecology processes. Detection of the DFE marginalized to an individual phenotype of interest, particularly beyond targeted segments such as coding regions, then can be informative to adaptation, mutational load, mutational target, and lethality. Such insight is relevant to elucidating the manner and speed with which evolution proceeds, predicting the trajectory of future variants, and comparing traits, independently structured lineages, and environmental conditions.

In contrast to tools that require the same individual-level genotypes and phenotypes employed as input for a GWAS, many techniques now typically take advantage of the summary statistics resulting from a GWAS that often are already publicly available. This provides much greater accessibility and convenience, not the least of which a substantial decrease in computational expense [18]. Notably, some methods indirectly utilize GWAS summary statistics to stringently subset the input data, thereby discarding the vast majority of information [19,20], but a much more desirable alternative would be to explicitly incorporate an entire set of genetic markers with a joint probabilistic model that unites evolutionary processes with genomic architecture [5,21,22]. A promising avenue to achieve this objective is the Poisson Random Field (PRF), which uses diffusion approximation to model allele counts for a given sample size conditional on parameterizations of demography and selection [23–26]. This calculation yields an expected site frequency spectrum that can be treated as a probability distribution for independent SNP data, as has been previously done to estimate a generalized DFE among coding regions [27]. Importantly, the assumption of independence between loci consequently does not address the influence of linkage disequilibrium (LD). However, while it would be ideal to explicitly model the full relationship among markers, such an endeavor would be too computationally intensive for practical implementation. Conversely, the PRF acts as a useful yet principled approximation by ignoring LD and thus allowing a composite likelihood across sites while still permitting maximum likelihood estimation of relevant parameters. This composite likelihood approach offers the large benefit of exploiting genomic-scale data efficiently, including integrating with a simple and computationally inexpensive model of GWAS summary statistics and true effect sizes. Additionally, the PRF is optimized for very weak selection coefficients, particularly at a scale much lower than typically explored for investigations of this nature.

Motivated by this potential to obtain a genome-wide DFE from modeling SNP-specific selection coefficients with the PRF, we present ASSESS (Association Summary Statistics for Estimating Selection among Sites) as a Python2 module for inferring trait-specific fitness effects from observed genome-wide allele sample frequency and GWAS summary statistic data per SNP. In this article, we introduce our likelihood-based model, represent its power and robustness through various *in silico* validations, and further illustrate its proof-of-concept with an empirical investigation. Importantly, we exhibit the ability of ASSESS to retain accuracy under several cases of model misspecification, including LD causing correlation structures within both allele count and estimated effect size input data vectors, and assumption violations of the genomic architecture. Subsequently, we demonstrate ASSESS usage on open-access GWAS datasets derived from the UK Biobank. These analyses exemplify the promising potential of ASSESS to obtain greater understanding of how natural selection regulates highly polygenic quantitative traits and disease.

## Results

### ASSESS joins population genetic theory with a quantitative model of genomic architecture to estimate a DFE corresponding to a complex trait (model description)

ASSESS directly captures a trait-specific DFE by deploying the PRF to model selection coefficients against sample allele counts, while simultaneously leveraging GWAS summary statistics to inform *β*_*i*_, the true effect size on a particular phenotype by an individual SNP *i* (Figure 1). The input, which favorably is sourced from only two dataset types that generally are easily accessible, is exploited to infer three parameters: 1) *ω*_0_, a weighted point-mass on zero that is informed by a Laplacian prior; 2) *σ*, standard deviation for a Gaussian normal distribution with a mean of zero; and 3) *c*, a genome-wide constant that governs the linear relationship between the DFE and true effect size (2*N*_*e*_*s*_*i*_ = *c*|*β*_*i*_|) [28]. The quantities of *ω*_0_ and *σ* comprise a mixture model for *β*_*i*_ [29–31], which acts as a latent variable and thus is numerically integrated within the likelihood equation. Assuming fitness consequences are entirely dependent on the impact of a genetic site onto a trait, is then a scalar that transforms the genome-wide distribution of *β*_*i*_ into population-scaled selection coefficients. Notably, the sign for *c* indicates either positive or negative selection ubiquitously among analyzed polymorphisms (*i*.*e*. larger phenotypic effects, regardless of directionality, translate to stronger fitness effects, which are exclusively beneficial or purifying for a given dataset); here, we focus solely on negative following the rationale that new mutations are deleterious when stabilizing selection acts on a polygenic trait, which we perceive to be the most conventional scenario. As a result of this simplified structure for the DFE, *ω*_0_ signifies the proportion of loci that are of zero consequence to the phenotype in both functional effect and fitness, while 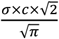 represents the expected selection strength across non-neutral markers given that the distribution of *β*_*i*_ is folded into the half-normal distribution of the DFE.

**Figure 1.**
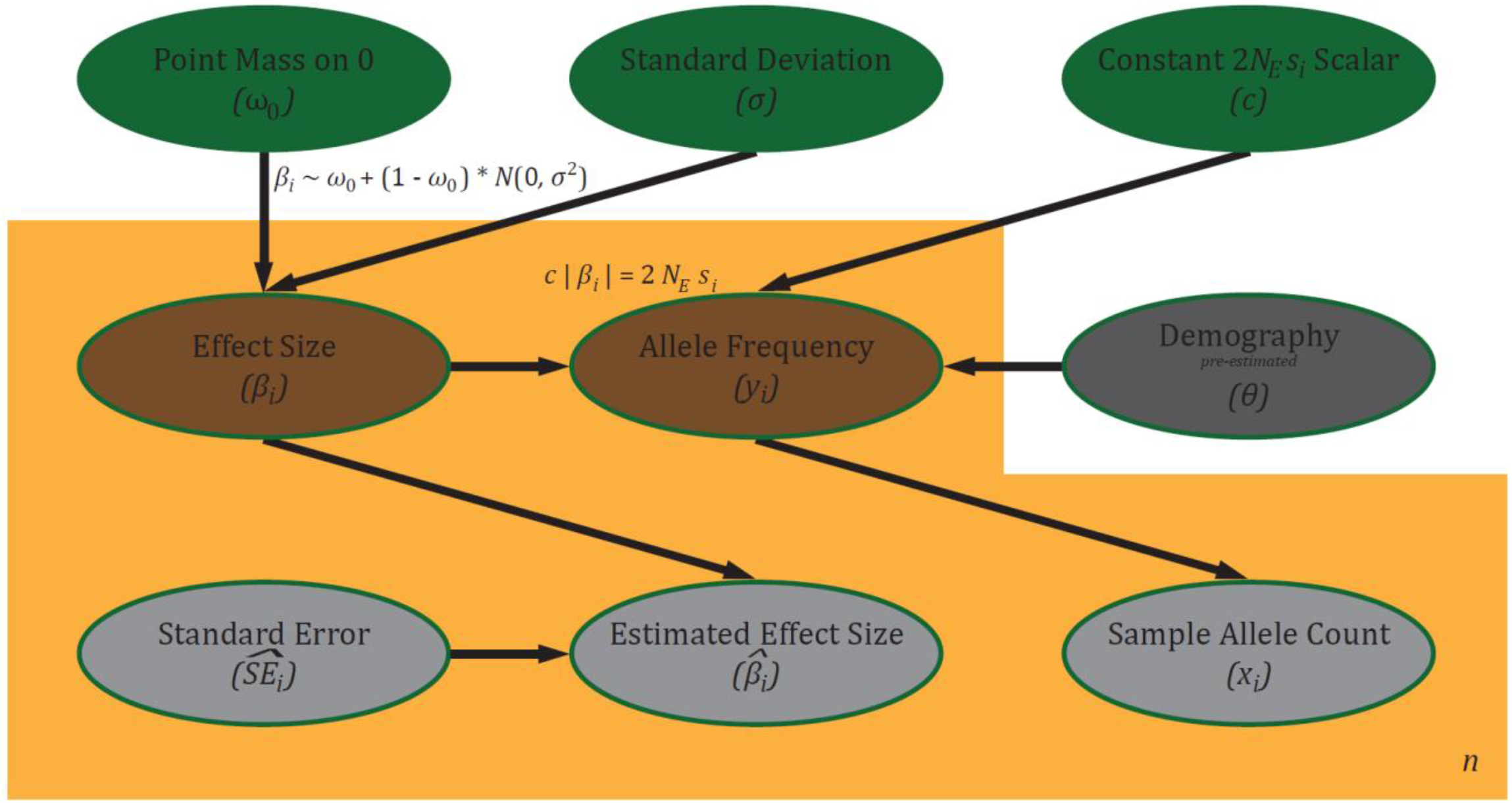
Probabilistic graphical model of ASSESS inferential framework. Free parameters of interest are in green, latent variables are in brown, and observed values are in gray (with demography “observed” in the sense that it is pre-estimated). The proportion of functional sites is controlled by the mixture component *ω*_0_, with non-zero true effect sizes (*β*_*i*_) modeled by a normal distribution centered on zero and standard deviation parameterized by *σ*. The GWAS summary statistic 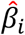 is then, assuming a normal distribution, informed by *β*_*i*_, which is numerically integrated, along with the GWAS-derived 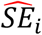. Allele count (*x*_*i*_) is conditional on the population-scaled selection coefficient, which is converted from *β*_*i*_ via *c*, under the PRF with demographic specifications separately inferred against the data, generically notated here as θ. Notably, due to the direct relationship between selection and effect size,*c* is irrelevant for SNPs with zero effect on the trait of interest. Additionally, usage of the PRF here allows integration of the true population-level allele frequency (*y*_*i*_).

### ASSESS can robustly recover the true DFE

To test the performance of ASSESS, we conducted *in silico* experiments against simulated DNA sequences and GWAS summary statistics (Table S1). We find no bias in the median DFE inferred amid 100 datasets (Figure 2). There is noticeable variance across the estimates, which is driven especially from a few outliers. However, this is within the context of an extremely high resolution in selection magnitude, *i*.*e*. | 2*N*_*e*_*s*_*i*_ | < 2.0, with most error occurring in the weakest bin of | 2*N*_*e*_*s*_*i*_ | < 2.0. Evaluation against additional simulation sets (Table S1) reveal that these favorable results are largely maintained regardless of: tuning parameterization (Figure S1); genomic processes such as recombination rate, mutation rate, and coefficient of allele dominance (Figure S2); sampling of individuals for both allele counts and GWAS summary statistics (Figure S3); assumption violations in how the latent true effect size is obtained (Figure S4); and single-population instantaneous size changes across three discrete epochs (Figure S5). Particularly notable is that simulations that challenge our assumption of a direct linear relationship between selection and effect size, including incurring decreased heritability and variance due to environmental effects, reflected no noticeable difference in the results (Figures S4 – S5). Likewise, uncertainty in the demographic background appeared to have no impact on the analysis (Figure S5). Together, these exercises demonstrate consistent behavior in uncovering the DFE throughout an array of conditions, suggesting the promise of ASSESS to reach valuable conclusions when exposed to real data.

**Figure 2.**
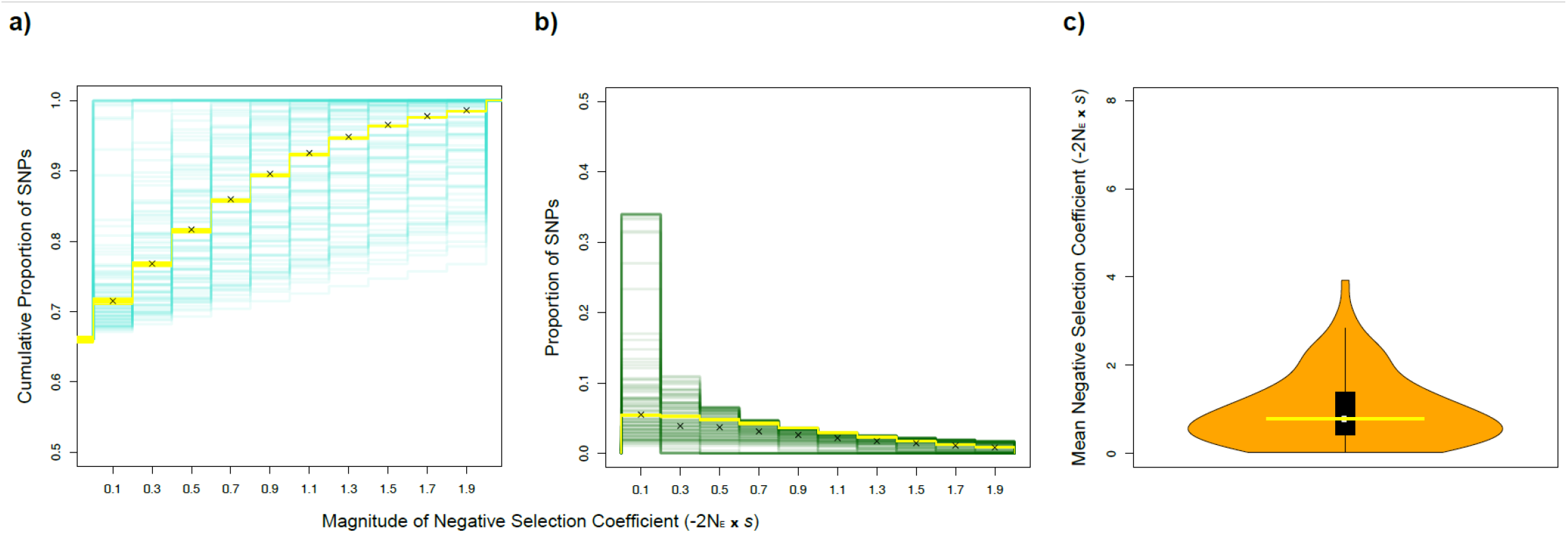
ASSESS performance given a simulated history of constant population size. **a, b)** Yellow lines indicate true values while teal/green lines represent the associated independent inferences of the DFE among 100 simulated datasets, with black marks denoting the median estimate. The x-axis, which covers a range of very weak selection coefficients, is presented in discretized positive units of increasing selection strength (*i*.*e*. scale of −2*N*_*e*_*s*_*i*_) for visual convenience. **a)** The y-axis plots the cumulative density of SNPs, normalized as a proportion of the total set including sites with no functional effect as well as loci undergoing strong selection. **b)** The y-axis plots the DFE, normalized as a proportion of the total set including sites with no functional effect as well as loci undergoing strong selection. **c)** Yellow boxplot indicates true values while orange violin plot and embedded black boxplot represent inferences of the mean average for the functional component of the DFE (presented in positive units, *i*.*e*. scale of −2*N*_*e*_*s*_*i*_). The range of the y-axis corresponds to the total optimization search space.

### ASSESS maintains accuracy in the face of severe genomic architecture misspecification

To further challenge the robustness of ASSESS, we included a set of simulations that incorporated additional assumption violations. Specifically, the underlying distribution of effect sizes was governed by an exponential distribution rather than a Gaussian normal, which is also an additional stress to our modeling of selection and effect size (Table S1). Moreover, we performed a set of inferences wherein the informed prior on ω_0_ was misspecified. This experiment yielded some bias from the deployment of the exponential distribution (Figure S6), especially in comparison to the previous efforts. However, there is no change in the median error, and the overall variance has noticeably decreased, though the minimum error has also increased (Figure S6d). Additionally, ASSESS accommodated the *a priori* inaccuracy in ω_0_ in excellent fashion, with no visible difference in the estimates. This exercise, which combined a dynamic demographic history along with several ASSESS model misspecifications regarding LD, effect of selection on phenotype, and genomic architecture, provided a formidable test to demonstrate the potential utility of ASSESS for real data.

### ASSESS recovers a range of DFEs for UK BioBank Traits

For our empirical application, we contextualized our empirical analysis against the estimates produced by Zeng *et al*. (2021), which we find to be the most similar implementation to ASSESS. However, though Zeng *et al*. (2021) explored a parameter correlated with fitness effects, their inferred property is nonetheless fundamentally different, thus a direct quantitative comparison is not possible. To this end, we selected two traits, specifically one under very strong selection and one under very weak selection based on the Zeng *et al*. (2021) inference, for a qualitative comparison of rank order per each of four UK Biobank trait categories. In three of the four categories, our findings are in agreement regarding which trait is under strong or weak selection (Figure 3). For the category of physical measures, we also added BMI due to its historical comparisons with height (*e*.*g*. [12]), and likewise found congruence with Zeng *et al*. (2021), as well as the conventional thought, in its inferred fitness effects relative to height. Hence, while a precise comparison to previously published results is difficult, this approximation via ranks nevertheless suggests general concordance.

**Figure 3.**
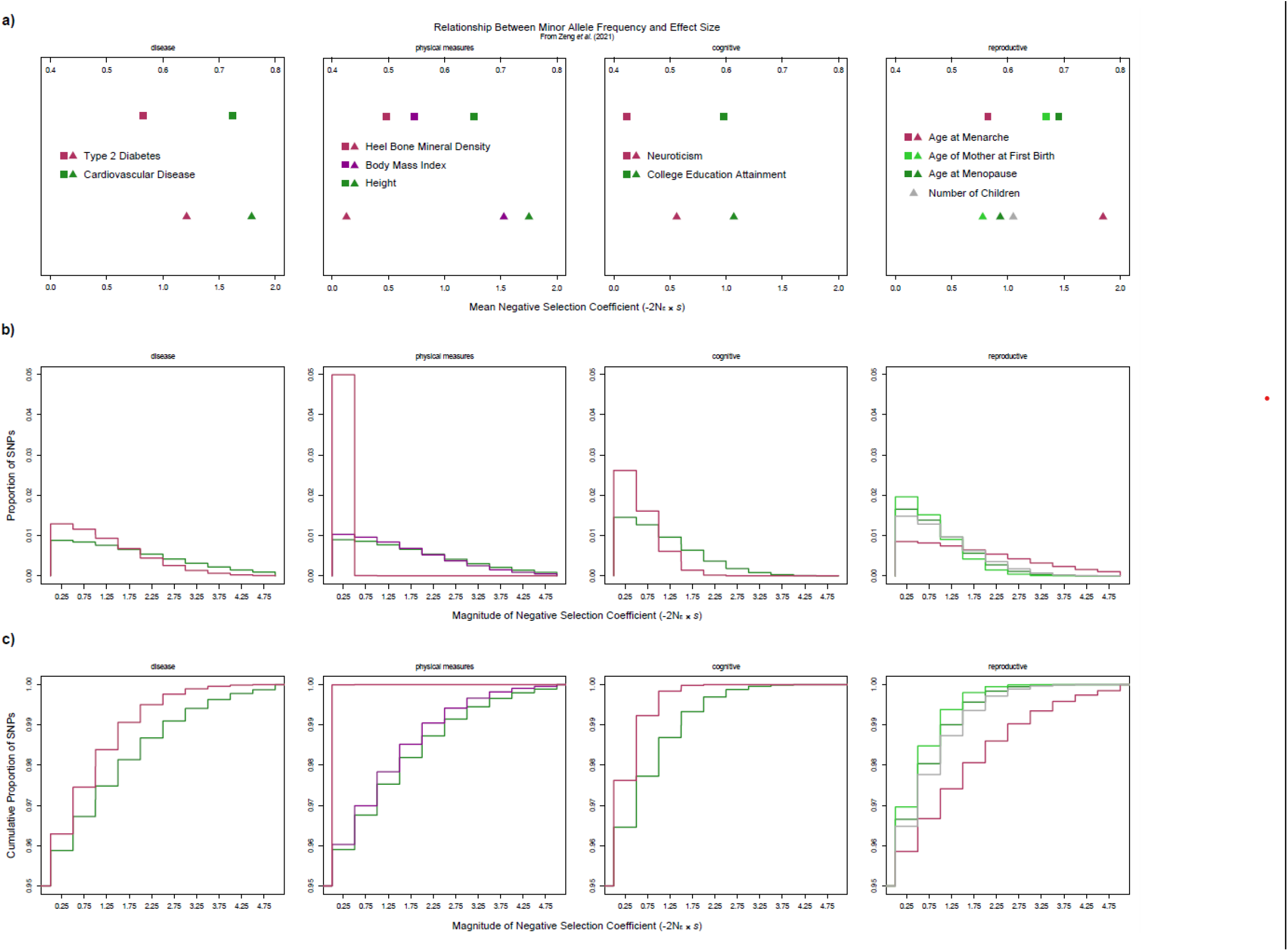
Selection inference for UK Biobank traits using ASSESS. Plots with the same x-axis unit have the same range among the four categories (*i*.*e*. the scaling remains the same horizontally across plots). **a)** The top half of the plots, which contain square data points, are estimates from Zeng *et al*. (2021), while the bottom half of the plots, which contain triangle data points, are corresponding empirical inferences from this study. Importantly, these two sets of results are of correlated yet distinctly different quantities; Zeng *et al*. (2021) investigated the relationship between minor allele frequency and effect size, whereas we focused on the expected value of the DFE (disregarding neutral sites). As a result, this is primarily a qualitative comparison, with the x-axis scale for the Zeng *et al*. (2021) and ASSESS estimates on the top and bottom, respectively. **b, c)** Color scheme for individual traits follow the legend in a). **b)** The y-axis plots the normalized DFE of the ASSESS empirical inferences. **c)** The y-axis plots the normalized cumulative density of SNPs of the ASSESS empirical inferences.

To better explore the inconsistency between the two studies for the category of reproductive phenotypes, we analyzed four datasets in total; this includes number of children, which although not part of Zeng *et al*. (2021), we decided to report since it is the most direct measurement of fitness. While the relative relationship between the estimates for first birth and menopause ages are quite similar, there is a strong disparity in the inference for age at menarche. However, it is important to consider that the parameter detected by Zeng *et al*. (2021) had a moderate correlation with selection strength, thus it is not expected to exactly reproduce a ranking of selection intensities. This is especially relevant here since the uncertainty estimated for the reproductive traits were overlapping in Zeng *et al*. (2021). Additionally, their methodology intended for a much different selection regime than is operated by ASSESS, with their study targeting selection coefficients up to three orders of magnitude greater than the resolution ASSESS is best suited (*i*.*e*. |2*N*_*e*_*s*_*i*_ | <2.0). Furthermore, the inherently complex nature of reproduction combined with its intimate ties to fitness possibly incurs greater sensitivity to methodological differences and thus produces much more variance in results. Interestingly, the fitness effects detected for number of children is quite moderate in magnitude (a very close approximation to the average among all traits), which perhaps exemplifies its high dimensionality to the point of effectively representing all traits simultaneously.

## Discussion

This study illustrates the potential of ASSESS to detect genome-wide selection coefficients associated with a complex polygenic trait of interest. Our *in silico* experiments demonstrate that ASSESS remains robust across a range of tuning parameterizations, data properties, and genomic architectures, including a plethora of flagrant model misspecifications. In particular, we discover that in spite of the strict linear relationship enforced between selection coefficient and effect size, ASSESS behavior is stable amid more dynamic simulation models whereby the true trait effect distribution has a likely more realistic transformation into the DFE. In particular, decreasing the genotype-phenotype correlation and level of heritability showcased the ability of ASSESS to tolerate extrinsic forces influencing trait expression. We posit that the 2*N*_*e*_*s*_*i*_ = *c*| *β*_*i*_| relationship allows the parameter to “absorb” various confounding factors that are not addressed by our model, thus the simple linear regression of the true selection coefficient against the true functional impact sufficiently captures the DFE from the observed data. This is perhaps supported by previous work that demonstrated decoupling between environmental effects and the DFE [32]. Moreover, this may also have assisted in resolving the differing distribution type for the effect sizes.

A major advantage of ASSESS is its usage of the PRF, which allows efficient computation due to its assumption of independent sites. However, this creates a major concern of the confounding effects from LD, which are inherently ignored by ASSESS due to this property of the PRF. Specifically, there are two avenues by which linkage can disrupt the underlying signal in the data: 1) the population genetic portion of the model – individual sites under selection induce an impact on allele frequencies for neighboring neutral SNPs, though negative selection should have a much less profound effect than a selective sweep signature; and 2) the genomic architecture portion of the model – non-causal genetic markers in close proximity to functional polymorphisms have an artificially inflated correlation with phenotypic values, thereby incurring error in the GWAS estimation of effect size. Fortunately, favorable conclusions from the simulation tests, all of which incorporated these two types of LD consequences, alleviate this factor. This is especially exemplified in the trials that varied recombination rates across a total span of two orders of magnitude. However, the inflated variance among replicates, especially within the weakest selection bin, may indeed be from the influence of linkage; this could be less problematic though for cases wherein the proportion of functional sites is relatively low.

Importantly, different combinations of parameter values can conceptually produce similar DFEs. For example, lowering *ω*_0_ could largely offset decreasing *σ*, and likewise reducing the intensity of the effect size architecture can be compensated by magnifying magnitudes of *c*. While the structure of the probabilistic model as informed by the allele counts and GWAS summary statistics should theoretically resolve these separate properties, including disentangling the effect size architecture from the DFE, the information may not be strong enough to tractably uncover these values; notably, this may be an avenue whereby LD has a particularly prominent effect. Indeed, we experienced preliminary difficulties in this regard, hence our informed prior with respect to *ω*_0_. While this is less than desirable, we found that the degrees of freedom had to be more limited, and we expect that polygenicity can be reasonably attained *a priori* for many datasets. Importantly, while individual estimates can be obtained for *σ* and *c*, these are probably not interpretable under the inferential framework of ASSESS due to its simplifying assumptions; as previously alluded, these parameters may be capturing unintended signals in service of ASSESS optimizing 2*N*_*e*_*s*_*i*_, thus are unreliable individually.

Interestingly, the quantity inferred by ASSESS deviates from a traditional perspective of the DFE. Our method of course has the feature of extracting a marginalized distribution, which is specified to a putative trait, from a theoretical aggregate of generalized fitness effects, which is a more commonplace construction of the DFE. Beyond this though, the modeling framework of ASSESS incurs additional atypical elements. First, whereas the target of obtaining the DFE is usually confined to a genomic subset, ASSESS is designed to be agnostic to type of genomic region and thus potentially genome-wide. However, the inference is ultimately dataset-dependent, thus wholly conditional on the site selection of the SNP chip that was used to generate GWAS data, which may not be entirely representational. Moreover, while ascertainment bias from allele frequency differences can be corrected within ASSESS, the impact of fixed mutations cannot be accommodated since our approach only operates on polymorphisms. As a result, the ASSESS DFE is partial to sites presently segregating within the collected data, therefore it cannot be interpreted as completely representing the predictive probability of generating fitness effects. Notably, while this elicits an omission of stronger negative selection coefficients, the focus on extremely small fitness consequences pairs well with the resolution of the PRF (*i*.*e*. 2N_e_s_i_ < 2.0). Our implementation then is able to discover a signature that can be challenging to capture on a highly polygenic scale and thereby may have been overlooked by other approaches. Interestingly, this heightened sensitivity to nuanced signals perhaps offers a compelling exploration of the genome under a more omnigenic perspective (*i*.*e*. one that considers contribution to a trait from a much greater mass of peripheral genes).

On that note, pleiotropy is another major consideration in the interpretation of our DFE. In particular, correlated traits would invoke a high overlap of the set of associated variants, thus ASSESS is potentially capturing a somewhat compounded DFE that describes several related traits. This begs the question of the exact definition of a trait, especially within the context of pleiotropy [11]. Theoretically, if the overall phenotype could be deconstructed into a suite of perfectly independent traits, then ASSESS is effectively aiming to discover the proportional contribution of each of these partitions to the absolute DFE. In practice though, traits are effectively an arbitrary artificial construction. To that end, a potentially interesting application of ASSESS then could be to compare estimated DFEs from seemingly related traits to reflect differences in pleiotropic effects. Similarly, inferences on the same trait from different populations could gain new insight for trait evolution.

A promising avenue to further develop this approach in a future implementation is to employ the simulation pipeline developed here coupled with a machine learning framework. This could allow a much greater level of complexity, such as incorporating pleiotropic interactions, environmental effects, positive selection with purifying selection, and temporal changes in phenotype optimum. Importantly, a simulation-based machine learning application could also possibly allow estimates of the individual parameters that define our DFE, including without a prior on the proportion of functional sites. These individual quantities can be of great interest with: 1) offering insight into mutational target size; 2) disentangling scenarios of increased polygenicity of weaker selection from decreased polygenicity of stronger selection; and 3) describing the relationship between selection and genomic architecture. Regardless, ASSESS demonstrates a promising and interesting application of the PRF to leverage GWAS summary statistics in a convenient and efficient manner for illuminating the genomic architecture of complex traits.

## Methods

### Likelihood-based Model

The baseline framework is a straightforward combination of the PRF for frequency changes of biallelic polymorphisms in response to selection and drift [23,24] and a sparse linear model for a complex trait that has been widely used in quantitative genetics [29–31] (Figure 1). These two components are linked by an assumed functional relationship between each site’s population-scaled selection coefficient, *S*_*i*_=2*N*_*e*_*s*_*i*_, and the corresponding true effect size, *β*_*i*_: *S*_*i*_ =*f*(*β*_*i*_; *c*)= *c* | *β*_*i*_ |, wherein *c* is a free parameter that controls the scale of the linear relationship [28].

For the observed allele count, *x*_*i*_, we deploy a standard PRF model that fits *S*_*i*_ conditional on a pre-estimated single-population demographic history with instantaneous change among discrete epochs of constant size. This approach implies binomial sampling of *x*_*i*_ given a true population-level allele frequency *y*_*i*_, which is integrated over. For the purposes of this paper, however, we treat the calculation of the PRF density function:

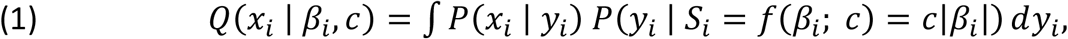

as a “black box” and execute it numerically given a discretization of 1,000 grid points using code borrowed from LASSIE [27]. For every possible *x*_*i*_ value, the density function *Q*(*x*_*i*_ | *β*_*i*_,*c*) is solved over a fine grid of *β*_*i*_ values and subsequently obtained by a table lookup per SNP. Notably, this calculation of *Q*(*x*_*i*_ | *β*_*i*_,*c*) allows for controlling uncertainty in the ancestral allele, akin to LASSIE. To address missing data, sampling level can subsequently be down-projected through the hypergeometric distribution [33,34].

To account for the GWAS process, we suppose that the resulting estimated effect size,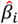, represents sampling from a Gaussian normal distribution whose mean equals the true value *β*_*i*_ [13,30,35] with standard deviation given by the estimated standard error,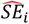:

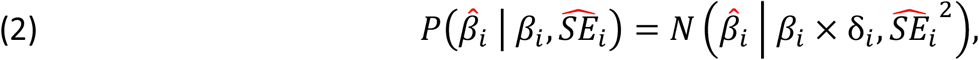

wherein *δ*_*i*_ is the standard deviation for the number of alternative alleles per sample in the case that 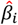 and 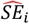 were obtained from standardized genotypes and thus *β*_*i*_ needs to be scaled proportionally (*δ*_*i*_ defaults to a value of 1 otherwise). We further employ a sparsity-inducing “spike and slab” prior distribution for the true *β*_*i*_, with a mixture coefficient for the weighted point-mass at zero, *ω*_0_, and variance, *σ*^2^, for the zero-centered Gaussian normal component:

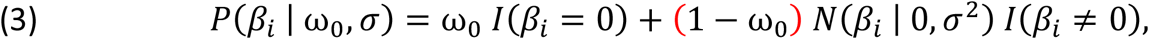

wherein *I* denotes an indicator function.

Combining these equations and assuming conditional independence of the population genetic data (*x*_*i*_), as in most applications of the PRF, as well as the quantitative genetic data (represented by the GWAS summary statistics 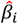 and 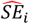) given the true value of_*βi*_, we obtain the likelihood function at a single locus *i*:

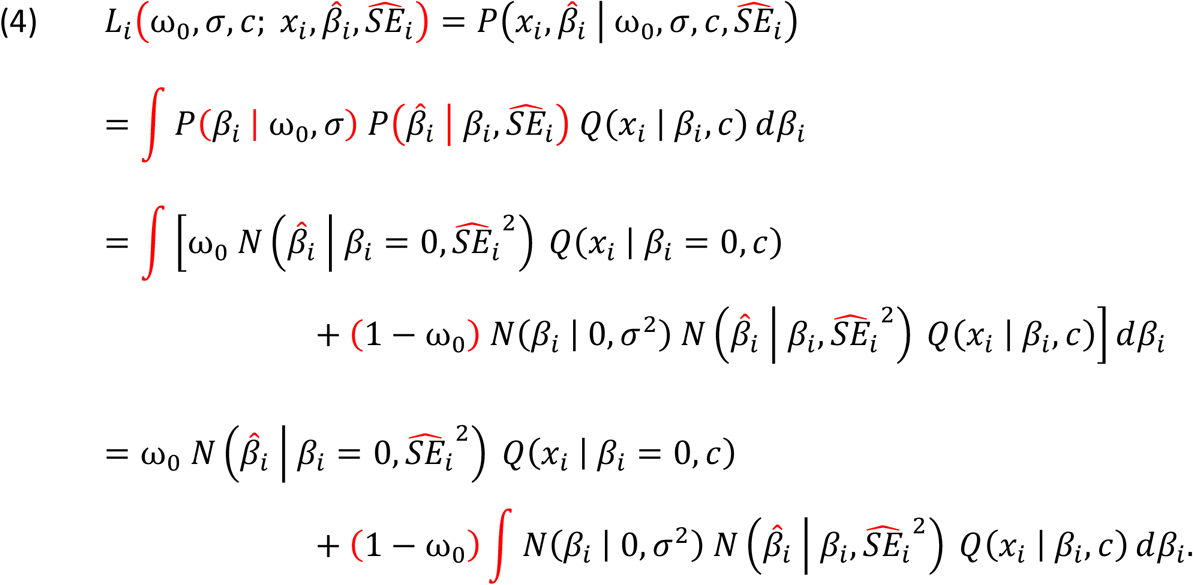

We approximate the integral over *β*_*i*_ numerically using the Gauss-Legendre quadrature rule, with nodes and weights scaled by 3× *σ*. A genome-wide set of markers, whereby *x*={*x*_*i*_} corresponds with 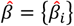 and 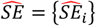, then yields the full likelihood function:

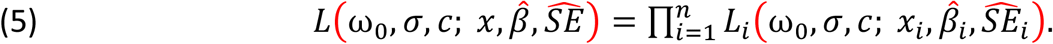

Therefore, the likelihood function has three total free parameters, two of which (ω_0_ and *σ*) define the prior distribution over the true effect size, and the third of which (*c*) defines the scale of the relationship between the true effect size and selection coefficient.

The commonly utilized platform to procure the genotypes considered in GWAS is the SNP chip, which tends to overrepresent variants segregating at higher minor allele frequency. Such ascertainment bias could be even further exacerbated by the SNP calling protocol or discordance in population structure between the samples informing the SNP chip design and GWAS individuals. To accommodate this, we make use of an importance weighting strategy. Here, *p*(*x*) represents the target distribution of relative frequencies over all possible minor allele counts given a reference panel, which for our empirical application is represented by complete genome sequences. Moreover, *Q*(*x*) represents the distribution of all possible minor allele counts for loci present within the GWAS data, upon which we are forced to operate despite our desire to exploit *p*(*x*) since our model depends on summary statistics.

Nevertheless, we can estimate the expected value for any function of interest, *f*(*x*), under the target distribution:

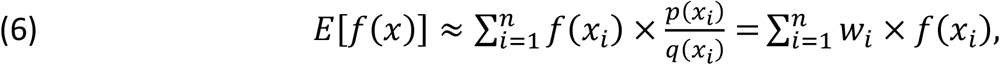

wherein 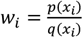 is derived for each possible minor allele count prior to optimization.

Therefore, casting the per-site log likelihood as *f*(*x*_*i*_), we obtain:

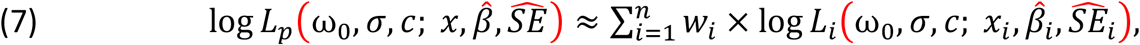

with down-projected through the hypergeometric distribution in cases of incomplete individual sampling, as done for *Q*(*x*_*i*_ | *β*_*i*_,*c*).

### Expectation-Maximization (EM) Algorithm

In the presence of observed values from*β*, the complete-data log likelihood function (CLL) for the baseline model can be expressed in terms of two sufficient statistics, *S*_0_ and *T*^2^:

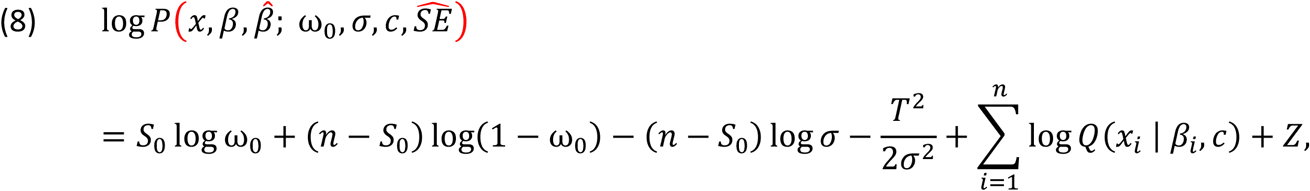

wherein *Z* is a quantity that does not depend on the free parameters. *S*_0_ represents the number of SNPs with effect sizes exactly equal to zero and *T*^2^ represents the sum of squares for the *β*_*i*_ values:

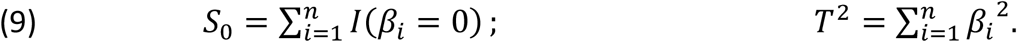

In this complete-data case, simple closed-form expressions are derived for maximum likelihood estimates (MLEs) of *ω*_0_ and *σ*^2^:

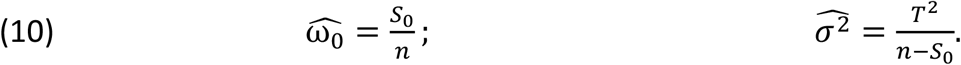

To curb potential identifiability issues stemming from the trade-off between these free parameters of our DFE construction, we deploy a Laplacian prior distribution on log *ω*_0_:

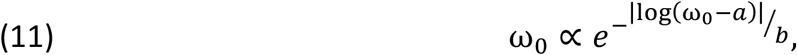

wherein *a* is the *a priori* expected value of log *ω*_0_ and *b* is a scale parameter positively related to the variance of *ω*_0_. This then transforms the CLL:

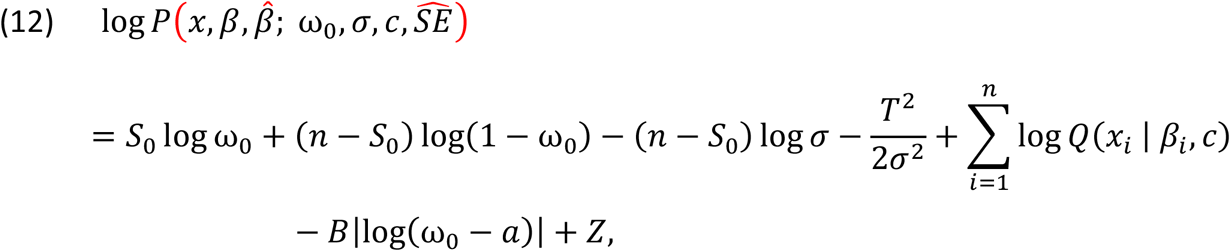

wherein 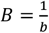, which acts as a penalty in space for departure from *a*.

In the usual way, an EM algorithm can be obtained by iteratively computing expected values of *S*_0_ and *T*^2^ (E step) and selecting values of *ω*_0_ and *σ*^2^ that maximize the expected CLL (M step). To achieve the expected values, Bayes’ rule is applied at each site:

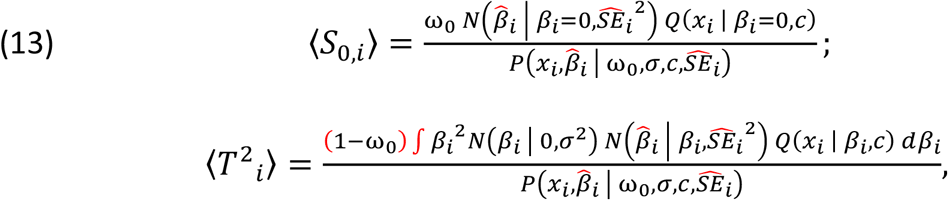

with all sites subsequently summed, such that 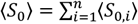 and 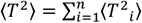; notably, the denominators here simply represent the per-site likelihood. For the MLE of *ω*_0_, due to our Laplacian prior, a modification must be employed in the following two cases:

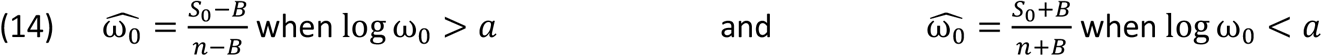

With all three calculations of 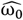 being potential maxima, given that *B* >0, then 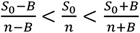 and therefore the global maximum is: 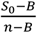 when 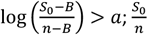 when 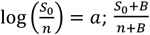 when 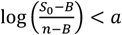 or whichever form of 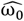 maximizes 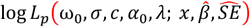 when the previous three conditions are not met. For implementation purposes, calculated values are forced to user-defined bounds when these are exceeded (typically 0.0 < *ω*_0_ < 1.0). To produce a MLE of *c*, which requires a numerical method, an update at each EM iteration is accomplished simply by a single step of gradient ascent, leading to a “generalized” EM algorithm.

### *In Silico* Experiments

To simulate test datasets, we developed a pipeline that exploits the software packages SLiM3 [36], msPrime [37], and simGWAS [38] to respectively generate DNA sequences, an initialized stable state of panmixia wherein only genetic drift occurs, and summary statistics. Specifically, SLiM3 simulated *y*_*i*_ values given a single-population history of either equilibrium or instantaneous size change across three epochs. For computational tractability, recombination and mutation rates were respectively set to 1.5*e* −7 and 1.25*e* −7 (excepting certain trials wherein one of these genomic properties was evaluated), which is one order of magnitude greater than accepted values for humans [39], with population size and temporal parameters correspondingly downscaled one order of magnitude from values relevant to human demography. Additionally, the coefficient of dominance equaled 0.5 for both neutral and selected alleles, barring individual tests wherein different values were tested (Figure S2; Table S1). To procure selection coefficients, which are specified as individual-level rates in SLiM3 versus population-scaled for ASSESS, draws were made from our “spike and slab” prior (except our single experiment utilizing an exponential distribution; Figure S6) assuming 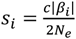 with the three free parameters pre-defined (Table S1) as well as *N*_*e*_=1,000 due to the aforementioned downscaling from human demography. Importantly for the three-epoch scenario, the true value for the population-scaled DFE (along with *σ* and *c*, regardless of generating value) is obscured since *s*_*i*_ is conditional on this reference *N*_*e*_ scalar, which represents a coarse approximation because of mutations randomly emerge throughout population size shifts over time. To address this, *S*_*i*_ was calculated from equally weighting *N*_0_ with the harmonic mean size during the trajectory of demographic change, hence creating a known DFE in the unit consistent with ASSESS estimates. For runtime efficiency, a single shared pool of 10,000 independent sequences equal in length was curated per experimental group (Table S1), from which there was a random subset of 1,000 to construct each of the 100 individual constituent datasets.

Afterward, msPrime recapitated neutral mutations segregating within a stable-size panmictic population prior to the emergence of a selected trait, thus allowing SLiM3 to efficiently bypass an incredibly long and resource-intensive burn-in period. With a complete genomic segment of population-level frequencies established, 100 diploid individuals (notwithstanding examinations of other sampling levels; Figure S3; Table S1) were randomly chosen to elicit *x*_*i*_ values, with monomorphisms pruned from the data and derived states beyond the oldest allele present coded the same so as to follow infinite sites. Notably, this simulation effort caused *ω*_0_, *σ*, and *n* to be governed stochastically, and in fact in a directional fashion from the *a priori* input values due to selection intensities eliciting differential fixation rates (*i*.*e*. mutations with stronger negative selection are more likely to be lost, thus inflating *ω*_0_ and deflating *σ ω*_0_ also drastically increases simply from the neutral sites introduced by this recapitation procedure). Consequently, true values could only be retrieved *post-hoc*.

Values for *β*_*i*_ were then calculated under two alternative models for the relationship between selection and effect size (Figures S4 – S6; Table S1), yielding two distinct datasets (though sharing identical allele count data). The first is based on the BayesS method, wherein *β*_*i*_ is calculated from the population-level allele frequency and this correlation is parameterized, thereby phenotypic contributions are naïve to selection coefficients (at least explicitly) but account for an environmental role [12]. Here, we simplified two of the parameters to be more aligned with ASSESS for the purpose of simulation efficiency: 1) the relationship between the variance of SNP effects and allele frequency was fixed to a constant specified *a priori* rather than randomly drawn per site, akin to the parameterization of gene-trait association from [40]; and 2) the common variance factor was set to a value such that the variance for the Gaussian component, given the mean of allele counts for markers under selection throughout the dataset, was equal to our *a priori* specification for *σ*^2^. The second strategy follows the seminal framework presented in [40] as modified by [41], wherein *β*_*i*_ derives from a more complex process that also incorporates heritability alongside all the variables already utilized in ASSESS and BayesS (*i*.*e*. fitness, allele frequency, and coupling between genetic variation and trait value). Notably, neither of these permit an exact equivalency for *c*, and likewise incur a different interpretation for *σ*, due to assumption differences from ASSESS (which are further exacerbated by linkage).

The final stage of this procedure involved simGWAS assigning summary statistics conditional on*β,x*, and GWAS sample size specifications (Table S1). Importantly, this approach considers covariance effects on 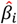 between adjacent loci, thus accounting for the influence of LD on the GWAS estimation process. However, this entailed computational restrictions, which was resolved by employing a sliding window with internal boundaries at every fourth polymorphism under selection from the first (*e*.*g*. 5^th^, 9^th^, 13^th^, etc.) up to the penultimate selected site per chromosomal segment. This paradigm allowed for overlapping sections, the absence of which would omit linkage dynamics at the edges, with at least two and up to five markers with functional effect for each simGWAS run.

### Empirical Application

Allele counts were retrieved from the 1000 Genomes phase 3 release while GWAS summary statistics, which were derived from the UK Biobank, were obtained from the lab website of Alkes Price [42]. This manner of data collection from two separate sources, wherein sample frequencies that are likely omitted from the GWAS study can instead be leveraged at higher resolution from an open-source repository containing many anonymized individuals across the whole genome, is what we envision to be most typical case. Given the drastic difference in amount of sites, non-intersecting loci between the two sets were culled and the discarded allele counts were used as part of the data for *a priori* demographic inference, a default feature of ASSESS. Notably, this independence in the data vector curation is not explicitly accounted for by the likelihood function, but this ought to be a rather minor consideration as long as the two data sources match in reference population.

### ASSESS Specifications

When implementing ASSESS, the underlying demography was correctly specified for all simulated scenarios apart from the single instance that explicitly investigated uncertainty in single-population size change history (Figure S5). Here, as well as for the entirety of the empirical application, epoch length (in units of 10,000 intervals with temporal length 1*e* −4) and relative *N*_*E*_ parameters were pre-estimated with a three-epoch instantaneous size change model utilizing LASSIE’s PRF implementation as called by ASSESS. This was executed against SNPs without GWAS summary statistics combined with polymorphisms within the lowest 10% quantile of 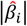 values. Aside from the *in silico* tests that directly stressed one of the following listed variables (Figure S1; Table S1), the tuning details for every inferential undertaking were as follows: search range for *σ* set automatically to 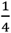 a log_10_ unit below and 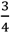 a log_10_ unit above the standard deviation of 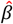 (the rationale for skewing the distribution higher in value is that a significant proportion of lower value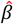 is expected to be captured by *ω*_0_); search range for *c* confined by the negative inverse of the upper bound for *σ* and 0.0 (thus the upper bound for the standard deviation of the functional component of the DFE, if it were hypothetically an unfolded normal distribution, is 1.0); nodes symmetrically centered at zero, as governed by the Gauss-Legendre quadrature rule, for numerical integration of *β*_*i*_; step size scalar of 1.0e −0.6 for gradient ascent of *c*; tolerance level of 2.220446049250313e − 09, for which when improvement of log ℒ is not greater than, optimization concludes and parameter values are estimated according to the local optimum; 50 maximum iterations of calculating for simulated data, and 10 maximum iterations of calculating for log ℒ empirical data, at which point optimization concludes and parameter values are estimated according to the local optimum; and 5 independent replicates of optimization cycles to approach a global maximum. For the simulation experiments, the Laplacian prior expected value for *ω*_0_ was always set to the correct value except for the single instance that tested this assumption (Figure S6; Table S1); for the empirical application, this was set to 0.95 in the interest of approximating the polygenicity detected from Zeng et al. (2021).

## Figure Legend

**Figure S1.**
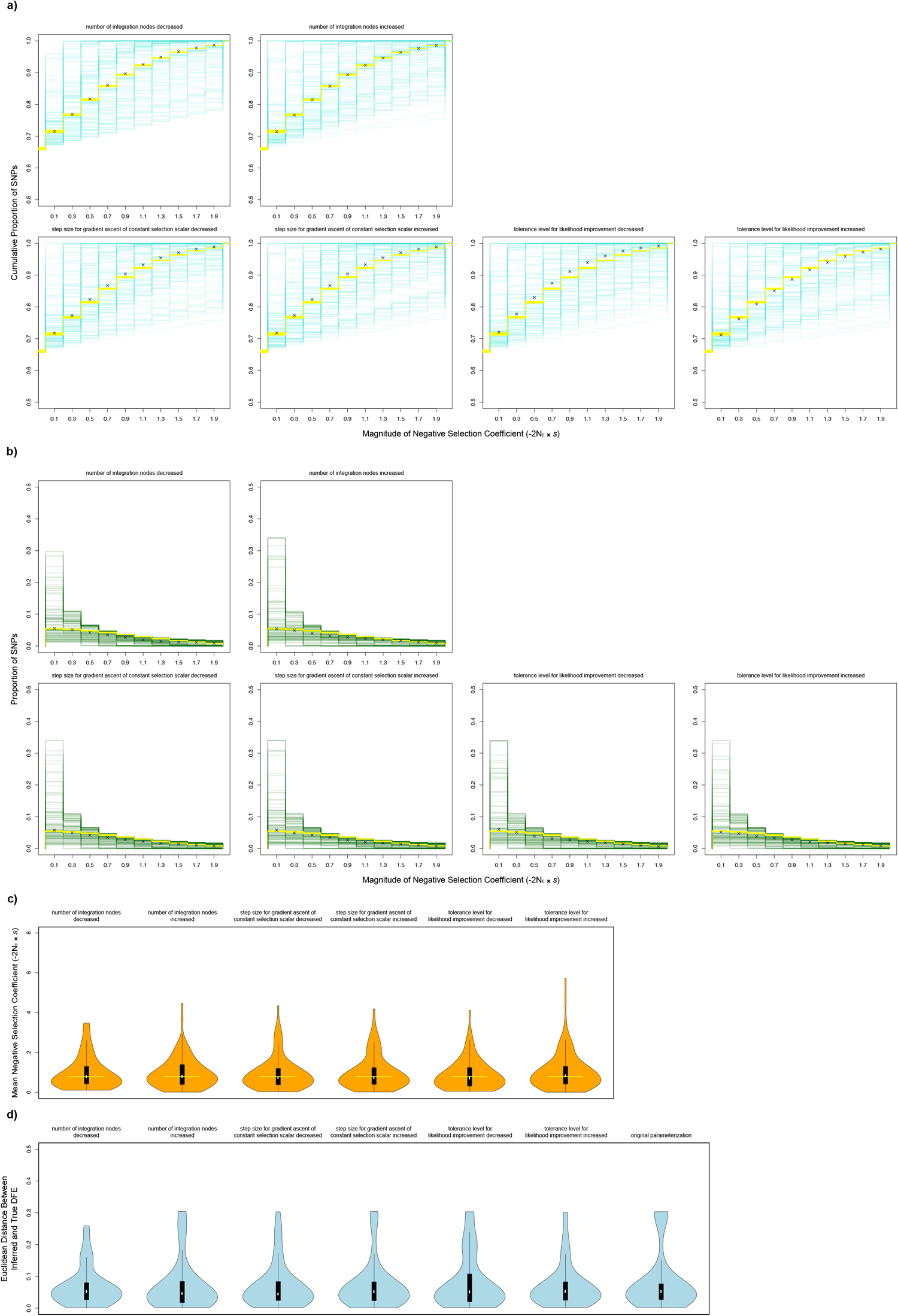
ASSESS performance across a range of optimization tuning parameterizations. The pictorial representation follows the same legend/structure as Figure 2. **d)** Blue violin plot and embedded black boxplot represent the Euclidean distance between estimated and true DFE, a goodness-of-fit metric that allows differences in overall sensitivity to be easily observed. The inferential application from Figure 2 is placed here as well for comparison.

**Figure S2.**
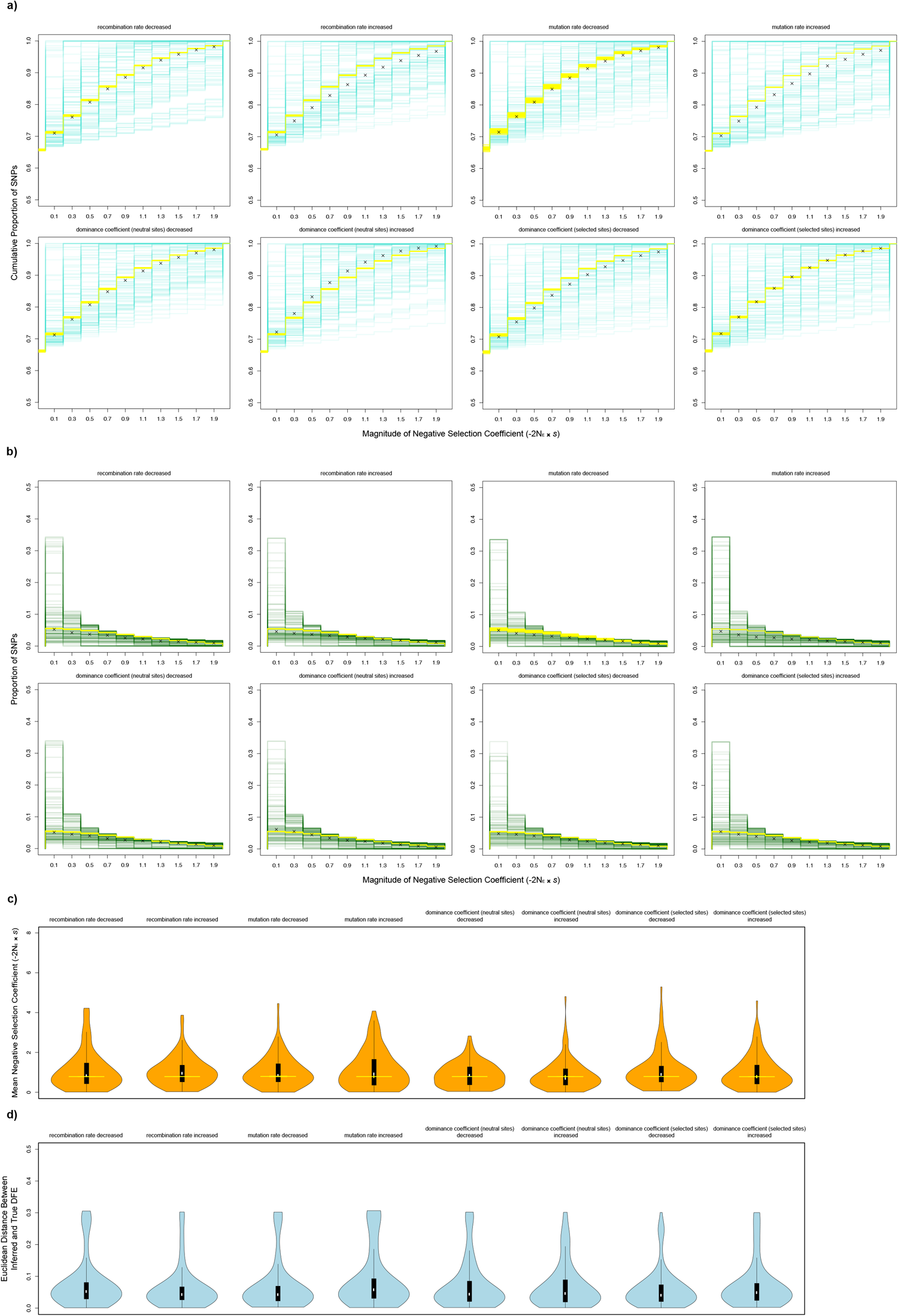
ASSESS performance across a range of genomic parameterizations. The pictorial representation follows the same legend/structure as Figure S1.

**Figure S3.**
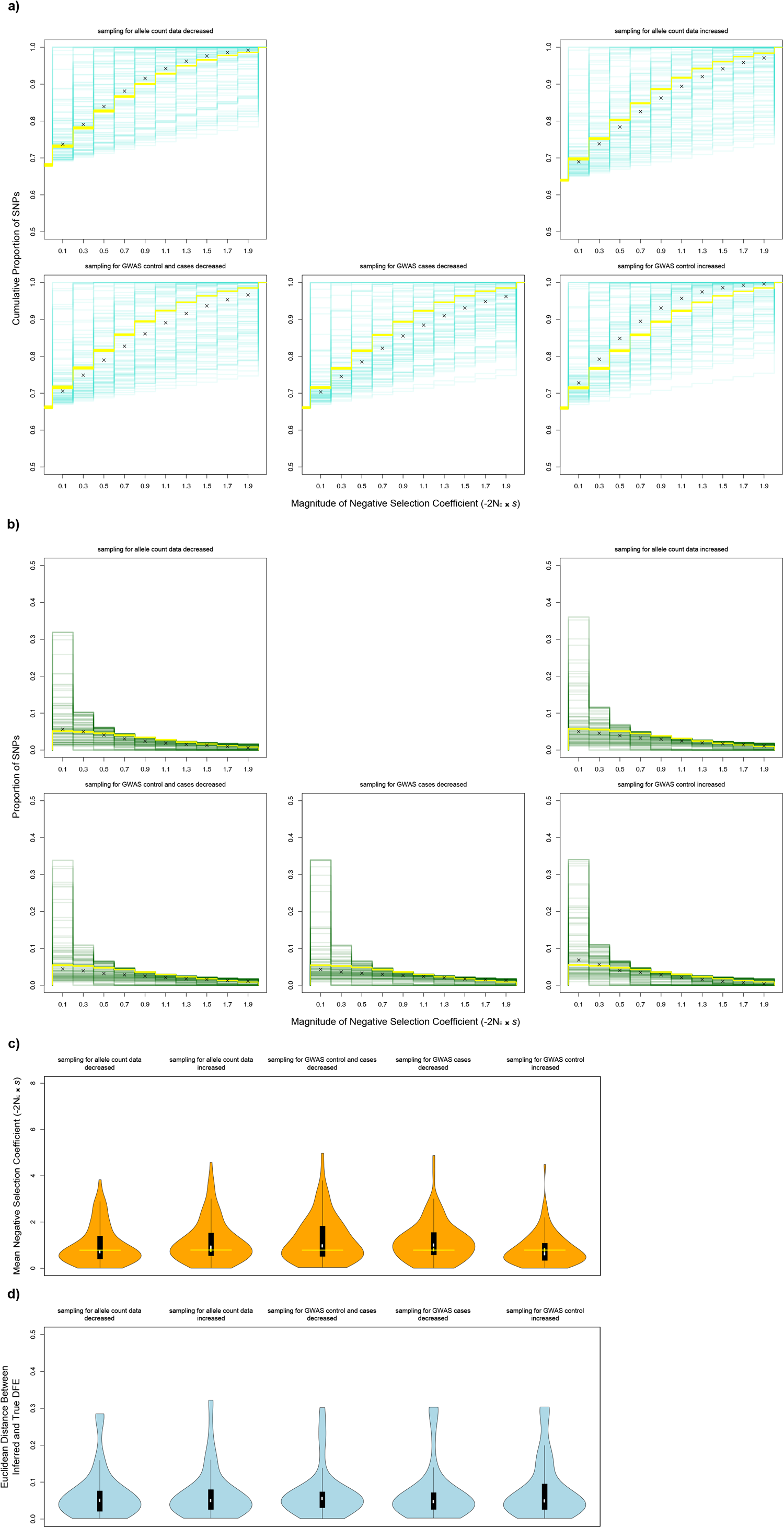
ASSESS performance across a range of sampling regimes. The pictorial representation follows the same legend/structure as Figure S1.

**Figure S4.**
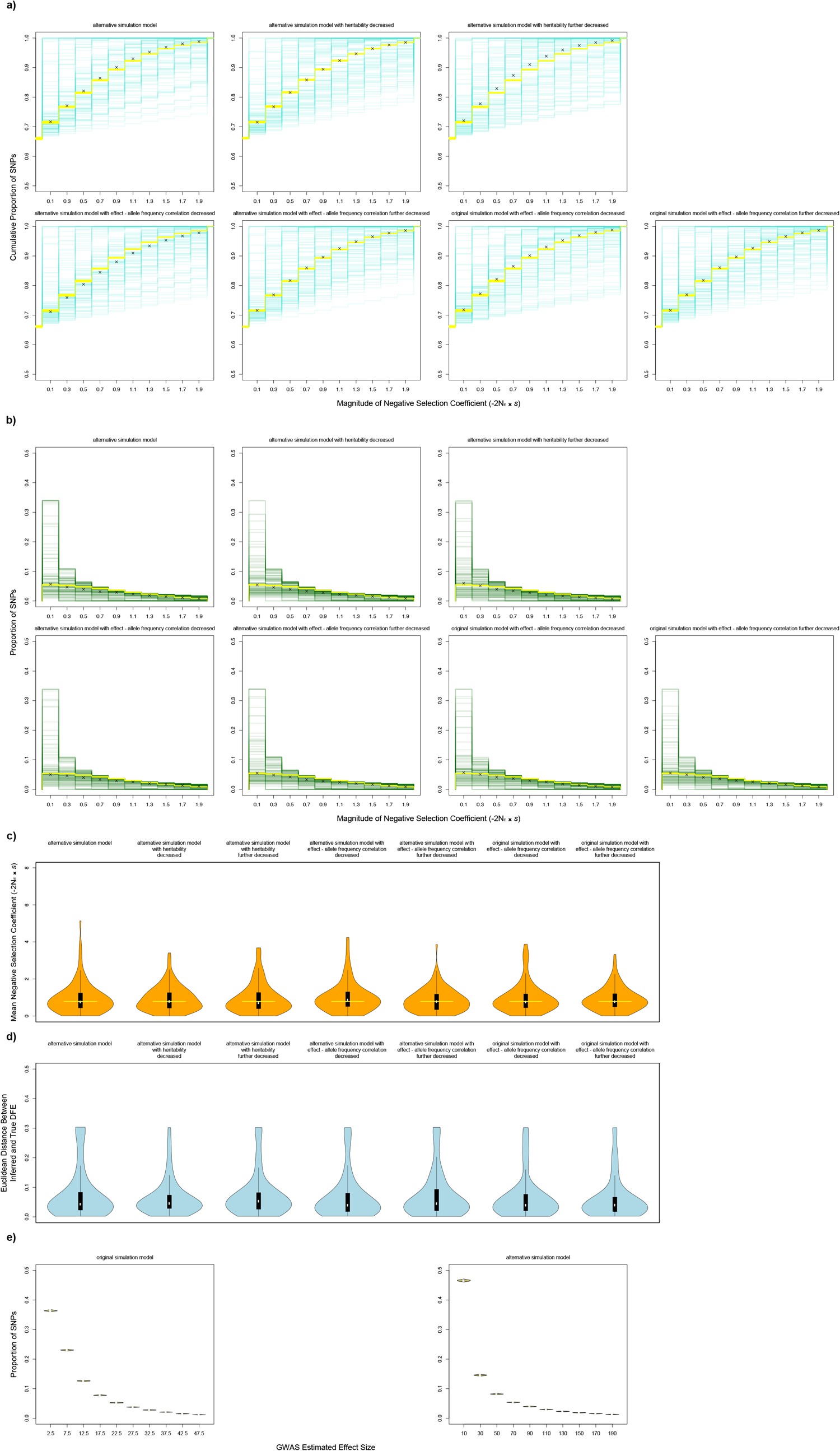
ASSESS performance across SNP effect parameterizations. Importantly, the alternative simulation model included here is much more highly parameterized and thus better accommodates realistic data. Moreover, added here are further analyses that varied generating values conferring a non-genetic impact, *i*.*e*. degree of heritability and extent that allele frequency corresponds to phenotype (Table S1). The continued success accomplished here validates ASSESS being agnostic to environmental effects and generally robust to model misspecification. The pictorial representation follows the same legend/structure as Figure S1. **e)** Yellow violin plot represents the distribution for simulated genome-wide estimated effect sizes, which includes non-functional loci. The x-axis is presented in discretized absolute value units, while the y-axis plots the proportion of SNPs from the total set of sites. Importantly, the disparity between the plots, particularly the distinctive distribution shapes on substantially different x-axis scales, depicts integrally different underlying functional relationships between selection and functional effect, providing a strong testing ground for ASSESS assumption violations.

**Figure S5.**
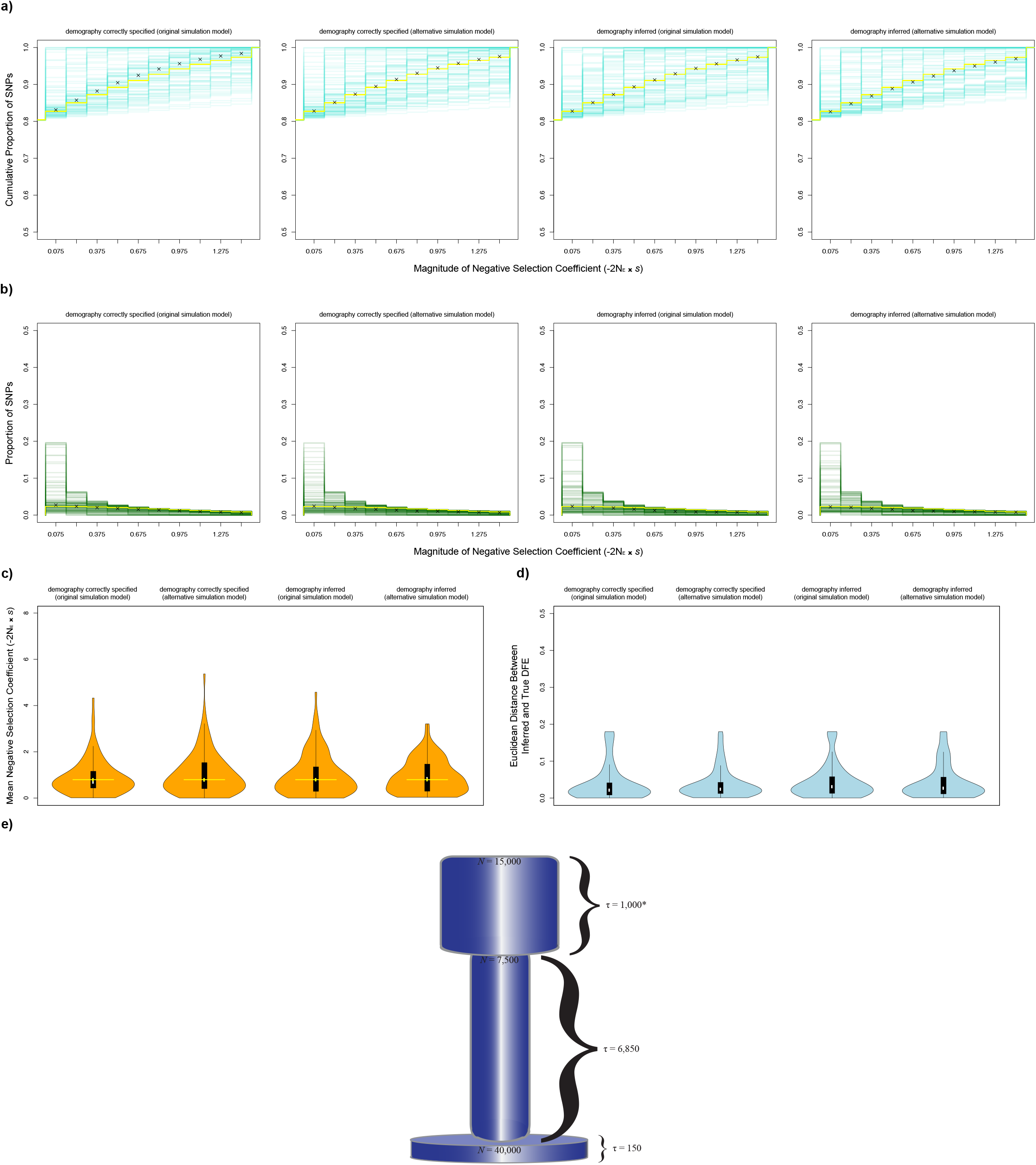
ASSESS performance given a simulated history of size change. Importantly, in addition to employing a scenario of demographic shifts, this dataset also utilized alternative DFE parameter values (notably at a decreased overall intensity) and GWAS sampling levels (Table S1). Estimates were performed under both the original and alternative simulation models. Moreover, included here is an additional inferential effort wherein demography was not correctly configured and instead pre-estimated from a subset of the data. Notably, due to differences in parameterizing the selection coefficient between ASSESS and the simulator (population-scaled versus rate, respectively), there was ambiguity over how to reconcile estimates with true values against a non-equilibrium population size, hence previous simulation experiments utilizing a constant size to avoid this artifact. An explanation regarding how the population size scaling factor was derived is in the Methods. The pictorial representation follows the same legend/structure as Figure S1. **e)** This is the assumed demographic model prior to rescaling, with units in diploid individuals and number of generations. **\*** The first epoch is generically set to 100 generations after rescaling, with a deeper neutral coalescent history accommodated by msPrime recapitation. To compare, the constant size history assumed 10,000 diploid individuals for 5,000 generations prior to rescaling and recapitation.

**Figure S6.**
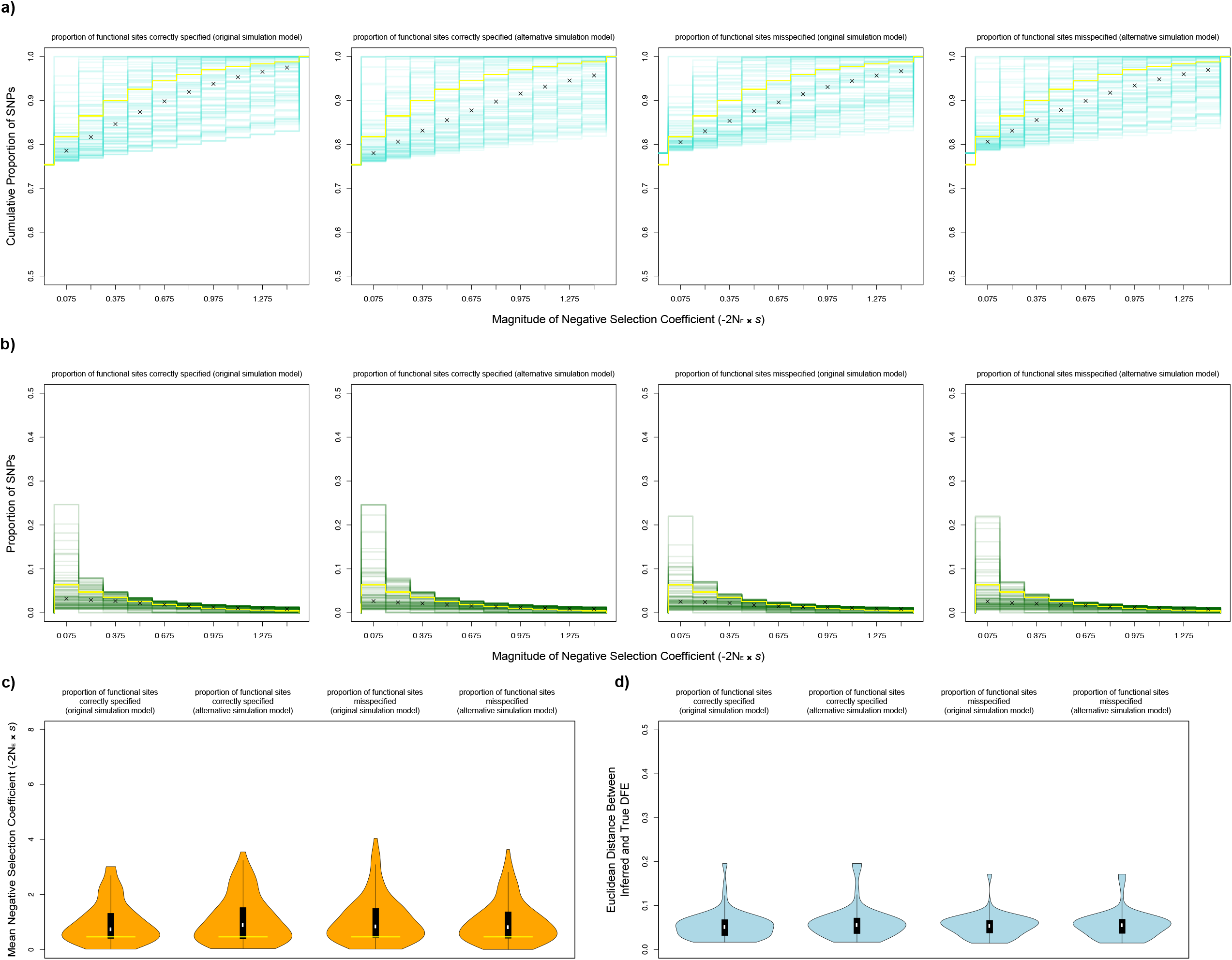
ASSESS robustness given misspecification of the DFE. This investigation, which utilized the demographic history of fluctuating population size as previously employed, challenged the genomic architecture underlying ASSESS by governing the simulated selection coefficients with a exponential distribution (Table S1). Estimates were performed under both the original and alternative simulation models, which (especially with the latter case) in conjunction with the altered distribution type, also violates the ASSESS assumption of a linear functional relationship between effect sizes and the DFE. Furthermore, inferences were additionally made with an incorrect prior for the point mass on zero. The pictorial representation follows the same legend/structure as Figure S1.

---

**Table S1.** Specifications for *In Silico* Experiments.

|  |  |  |
| --- | --- | --- |
| <u>original application</u><br>(Figure 1) |  |  |
| $\omega_0 = 0.05$ | $\sigma = 0.1$ | $c = -10.0$ |
| length of each independent sequence<br>= 25 KB | gene-trait coupling = 1.0 |  |
| sample size for GWAS control = 5,000 | sample size for GWAS case = 5,000 |  |
| *each additional simulated dataset for Figures S2 – S4 follows these same specifications*<br>*except when stated otherwise* |  |  |
| <u>experiment on resolution of numerical integration in ASSESS</u><br>(Figure S1) |  |  |
| additional inferential application with<br>decreased number of $\beta_i$ nodes = 20 | additional inferential application with<br>increased number of $\beta_i$ nodes = 100 | |
| <u>experiment on step size for gradient ascent of constant selection scalar in ASSESS</u><br>(Figure S1) |  |  |
| additional inferential application with<br>decreased step size scalar = $1.0e - 8$ | additional inferential application with<br>increased step size scalar = $1.0e - 4$ | |
| <u>experiment on tolerance level for log likelihood improvement in ASSESS</u><br>(Figure S1) |  |  |
| additional inferential application with<br>decreased likelihood tolerance = $1.0e - 13$ | additional inferential application with<br>increased likelihood tolerance = $1.0e - 5$ | |
| <u>experiment on recombination rate in simulations</u><br>(Figure S2) |  |  |
| additional dataset with decreased<br>recombination rate = $1.5e - 8$ | additional dataset with increased<br>recombination rate = $1.5e - 6$ | |
| <u>experiment on mutation rate in simulations</u><br>(Figure S2) |  |  |
| additional dataset with decreased<br>mutation rate = $1.25e - 8$ | additional dataset with increased<br>mutation rate = $1.25e - 6$ | |
| <u>experiment on coefficient of allele dominance in simulations</u><br>(Figure S2) |  |  |
| additional dataset with decreased<br>dominance coefficient for neutral sites = 0.2 | additional dataset with increased<br>dominance coefficient for neutral sites = 0.8 |  |
| additional dataset with decreased<br>dominance coefficient for selected sites = 0.2 | additional dataset with increased<br>dominance coefficient for selected sites = 0.8 |  |
| <u>experiment on sampling of individuals for allele count data in simulations</u><br>(Figure S3) |  |  |
| additional dataset with decreased<br>total allele counts = 100 (i.e. 50 diploids) | additional dataset with increased<br>total allele counts = 500 (i.e. 250 diploids) |  |
| <u>experiment on sampling of individuals for GWAS summary statistics in simulations</u><br>(Figure S3) |  |  |
| additional dataset with | additional dataset with | additional dataset with |
| decreased GWAS control<br>= 500 and cases = 500 | decreased GWAS<br>cases = 500 | increased GWAS<br>control = 50,000 |
| <u>experiment on alternative simulation model</u><br>*(Figure S4)* |  |  |
| additional dataset with<br>simulation model based on<br>Lohmueller (2014)<br>and heritability = 0.9 | additional dataset with<br>simulation model based on<br>Lohmueller (2014)<br>and heritability = 0.7 | additional dataset with<br>simulation model based on<br>Lohmueller (2014)<br>and heritability = 0.5 |
| additional dataset with simulation model<br>based on Lohmueller (2014),<br>heritability = 0.9,<br>and gene-trait coupling = 0.75 |  | additional dataset with simulation model<br>based on Lohmueller (2014),<br>heritability = 0.9,<br>and gene-trait coupling = 0.5 |
| additional dataset with<br>gene-trait coupling = 0.75 |  | additional dataset with<br>gene-trait coupling = 0.5 |
| <u>experiment on demographic history of instantaneous population size change</u><br>*(Figure S5)* |  |  |
| $\omega_0 = 0.2$ | $\sigma = 0.2$ | $c = -5.0$ |
| length of each independent<br>sequence = 250 KB | heritability = 0.3 | gene-trait coupling = 0.9 |
| sample size for GWAS control = 6,744 |  | sample size for GWAS case = 2,479 |
| <u>experiment on misspecified DFE shape in simulations and Laplacian prior of $\omega_0$ in ASSESS</u><br>*(Figure S6)* |  |  |
| $\lambda = 0.1$ | | $c = -5.0$ |
| common variance factor = 0.2 |  |  |
| length of each independent<br>sequence = 250 KB | heritability = 0.3 | gene-trait coupling = 0.9 |
| sample size for GWAS control = 6,744 |  | sample size for GWAS case = 2,479 |
| Laplacian prior of $\omega_0 = 0.753$ (versus 0.780) | | |

## References

1. Buniello A, Macarthur JAL, Cerezo M, Harris LW, Hayhurst J, Malangone C, et al. The NHGRI-EBI GWAS Catalog of published genome-wide association studies, targeted arrays and summary statistics 2019. Nucleic Acids Res. Oxford University Press; 2019;47: D1005–D1012. doi:10.1093/nar/gky1120

2. Visscher PM, Brown MA, McCarthy MI, Yang J. Five Years of GWAS Discovery. Am J Hum Genet. The American Society of Human Genetics; 2012;90: 7–24. doi:10.1016/j.ajhg.2011.11.029

3. Visscher PM, Wray NR, Zhang Q, Sklar P, McCarthy MI, Brown MA, et al. 10 Years of GWAS Discovery: Biology, Function, and Translation. Am J Hum Genet. Elsevier Company.; 2017;101: 5–22. doi:10.1016/j.ajhg.2017.06.005

4. Marjoram P, Zubair A, Nuzhdin S V. Post-GWAS: where next? More samples, more SNPs or more biology? Heredity (Edinb). Nature Publishing Group; 2014;112: 79–88. doi:10.1038/hdy.2013.52

5. Guo J, Yang J, Visscher PM. Leveraging GWAS for complex traits to detect signatures of natural selection in humans. Curr Opin Genet Dev. Elsevier Ltd; 2018;53: 9–14. doi:10.1016/j.gde.2018.05.012

6. Berg JJ, Coop G. A Population Genetic Signal of Polygenic Adaptation. PLoS Genet. 2014;10: e1004412. doi:10.1371/journal.pgen.1004412

7. Gazal S, Finucane HK, Furlotte NA, Loh P-R, Palamara PF, Liu X, et al. Linkage disequilibrium-dependent architecture of human complex traits shows action of negative selection. Nat Genet. Nature Publishing Group; 2017;49: 1421–1427. doi:10.1038/ng.3954

8. Abraham A, LaBella AL, Capra JA, Rokas A. Mosaic patterns of selection in genomic regions associated with diverse human traits. PLoS Genet. 2022;18: e1010494. doi:10.1371/journal.pgen.1010494

9. Gazal S, Loh P-R, Finucane HK, Ganna A, Schoech A, Sunyaev S, et al. Functional architecture of low-frequency variants highlights strength of negative selection across coding and non-coding annotations. Nat Genet. Springer US; 2018;50: 1600–1607. doi:10.1038/s41588-018-0231-8

10. Liu X, Loh P-R, O’Connor LJ, Gazal S, Schoech A, Maier RM, et al. Quantification of genetic components of population differentiation in UK Biobank traits reveals signals of polygenic selection. bioRxiv. 2018; 357483. doi:10.1101/357483

11. Simons YB, Bullaughey K, Hudson RR, Sella G. A population genetic interpretation of GWAS findings for human quantitative traits. PLoS Biol. 2018;16: e2002985. doi:10.1371/journal.pbio.2002985

12. Zeng J, de Vlaming R, Wu Y, Robinson MR, Lloyd-Jones LR, Yengo L, et al. Signatures of negative selection in the genetic architecture of human complex traits. Nat Genet. Springer US; 2018;50: 746–753. doi:10.1038/s41588-018-0101-4

13. O’Connor LJ, Schoech AP, Hormozdiari F, Gazal S, Patterson N, Price AL. Extreme Polygenicity of Complex Traits Is Explained by Negative Selection. Am J Hum Genet. Elsevier Company.; 2019;105: 456–476. doi:10.1016/j.ajhg.2019.07.003

14. Schoech AP, Jordan DM, Loh P-R, Gazal S, O’Connor LJ, Balick DJ, et al. Quantification of frequency-dependent genetic architectures in 25 UK Biobank traits reveals action of negative selection. Nat Commun. Springer US; 2019;10: 790. doi:10.1038/s41467-019-08424-6

15. Stern AJ, Speidel L, Zaitlen NA, Nielsen R. Disentangling selection on genetically correlated polygenic traits via whole-genome genealogies. Am J Hum Genet. Elsevier Company.; 2021;108: 219–239. doi:10.1016/j.ajhg.2020.12.005

16. Zeng J, Xue A, Jiang L, Lloyd-Jones LR, Wu Y, Wang H, et al. Widespread signatures of natural selection across human complex traits and functional genomic categories. Nat Commun. Springer US; 2021;12: 1164. doi:10.1038/s41467-021-21446-3

17. Eyre-Walker A, Keightley PD. The distribution of fitness effects of new mutations. Nat Rev Genet. 2007;8: 610–618. doi:10.1038/nrg2146

18. Pasaniuc B, Price AL. Dissecting the genetics of complex traits using summary association statistics. Nat Rev Genet. Nature Publishing Group; 2017;18: 117–127. doi:10.1038/nrg.2016.142

19. Yang J, Ferreira T, Morris AP, Medland SE, Madden PAF, Heath AC, et al. Conditional and joint multiple-SNP analysis of GWAS summary statistics identifies additional variants influencing complex traits. Nat Genet. Nature Publishing Group; 2012;44: 369–375. doi:10.1038/ng.2213

20. Field Y, Boyle EA, Telis N, Gao Z, Gaulton KJ, Golan D, et al. Detection of human adaptation during the past 2000 years. Science (80-). 2016;354: 760–764. doi:10.1126/science.aag0776

21. Price AL, Spencer CCA, Donnelly P. Progress and promise in understanding the genetic basis of common diseases. Proc R Soc B Biol Sci. 2015;282: 20151684. doi:10.1098/rspb.2015.1684

22. Uricchio LH. Evolutionary perspectives on polygenic selection, missing heritability, and GWAS. Hum Genet. 2020;139: 5–21. doi:10.1007/s00439-019-02040-6. Evolutionary

23. Kimura M. Diffusion Models in Population Genetics. J Appl Probab. 1964;1: 177–232. doi:10.2307/3211856

24. Sawyer SA, Hartl DL. Population Genetics of Polymorphism and Divergence. Genetics. 1992;132: 1161–1176.

25. Hernandez RD, Williamson SH, Bustamante CD. Context Dependence, Ancestral Misidentification, and Spurious Signatures of Natural Selection. Mol Biol Evol. 2007;24: 1792–1800. doi:10.1093/molbev/msm108

26. Gutenkunst RN, Hernandez RD, Williamson SH, Bustamante CD. Inferring the joint demographic history of multiple populations from multidimensional SNP frequency data. PLoS Genet. 2009;5: e1000695. doi:10.1371/journal.pgen.1000695

27. Huang Y-F, Siepel A. Estimation of allele-specific fitness effects across human protein-coding sequences and implications for disease. Genome Res. 2019;29: 1310–1321. doi:10.1101/gr.245522.118

28. Keightley PD, Hill WG. Variation maintained in quantitative traits with mutation-selection balance: pleiotropic side-effects on fitness traits. Proc R Soc B Biol Sci. 1990;242: 95–100. doi:10.1098/rspb.1990.0110

29. Zhou X, Carbonetto P, Stephens M. Polygenic Modeling with Bayesian Sparse Linear Mixed Models. PLoS Genet. 2013;9: e1003264. doi:10.1371/journal.pgen.1003264

30. Ma Y, Zhou X. Genetic prediction of complex traits with polygenic scores: a statistical review. Trends Genet. Elsevier Ltd; 2021;37: 995–1011. doi:10.1016/j.tig.2021.06.004

31. Zhao Z, Song J, Wang T, Lu Q. Polygenic risk scores: effect estimation and model optimization. Quant Biol. 2021;9: 133–140. doi:10.15302/j-qb-021-0238

32. Koch EM, Sunyaev SR. Maintenance of Complex Trait Variation: Classic Theory and Modern Data. Front Genet. 2021;12: 763363. doi:10.3389/fgene.2021.763363

33. Marth GT, Czabarka E, Murvai J, Sherry ST. The Allele Frequency Spectrum in Genome-Wide Human Variation Data Reveals Signals of Differential Demographic History in Three Large World Populations. Genetics. 2004;166: 351–372. doi:10.1534/genetics.166.1.351

34. Jouganous J, Long W, Ragsdale AP, Gravel S. Inferring the Joint Demographic History of Multiple Populations: Beyond the Diffusion Approximation. Genetics. 2017;206: 1549–1567. doi:10.1534/genetics.117.200493

35. Shi H, Kichaev G, Pasaniuc B. Contrasting the Genetic Architecture of 30 Complex Traits from Summary Association Data. Am J Hum Genet. American Society of Human Genetics; 2016;99: 139–153. doi:10.1016/j.ajhg.2016.05.013

36. Haller BC, Messer PW. SLiM 3: Forward genetic simulations beyond the Wright-Fisher model. bioRxiv. 2018; 418657. doi:10.1101/418657

37. Baumdicker F, Bisschop G, Goldstein D, Gower G, Ragsdale AP, Tsambos G, et al. Efficient ancestry and mutation simulation with msprime 1.0. Genetics. 2022;220: iyab229. doi:10.1093/genetics/iyab229

38. Fortune MD, Wallace C. simGWAS: a fast method for simulation of large scale case– control GWAS summary statistics. Bioinformatics. 2019;35: 1901–1906. doi:10.1093/bioinformatics/bty898

39. Schiffels S, Durbin R. Inferring human population size and separation history from multiple genome sequences. Nat Genet. Nature Publishing Group; 2014;46: 919–925. doi:10.1038/ng.3015

40. Eyre-Walker A. Genetic architecture of a complex trait and its implications for fitness and genome-wide association studies. Proc Natl Acad Sci. 2010;107: 1752–1756. doi:10.1073/pnas.0906182107

41. Lohmueller KE. The Impact of Population Demography and Selection on the Genetic Architecture of Complex Traits. PLoS Genet. 2014;10: e1004379. doi:10.1371/journal.pgen.1004379

42. Loh P, Kichaev G, Gazal S, Schoech AP, Price AL. Mixed-model association for biobank-scale datasets. Nat Genet. 2018;50: 906–908. doi:10.1038/s41588-018-0144-6

